# Vaginal lubricants in the couple trying-to-conceive: assessing healthcare professional recommendations and effect on in vitro sperm function

**DOI:** 10.1101/499293

**Authors:** S. C. Mackenzie, S. A. Gellatly

## Abstract

Vaginal lubricants are commonly used by couples trying-to-conceive. However, most vaginal lubricants are sperm toxic and therefore should not be used by couples trying-to-conceive. Despite this, lubricant sperm toxicity is insufficiently reported and guidance for healthcare professionals (HCPs) are absent. In this study, lubricant-related practices of fertility-based HCPs in Scotland were sampled via an online survey. Lubricants identified as being utilised in the fertility setting were subsequently incubated with prepared sperm samples to establish effects on sperm motility.

HCP recommendations (n=32) on lubricant use were varied although knowledge related to sperm toxicity was generally poor. HCPs infrequently asked about lubricant use and were unaware of guidance in this area. Aquagel, the only prescribed lubricant identified in this study, reduced sperm progressive motility to 49% of control after 10 minutes, even at concentrations as low as 5%. Vitality testing suggested the deterioration in progressive motility with Aquagel was not as a result of cell death. Conversely, Pré Vaginal Lubricant, a ‘sperm-safe’ lubricant, did not significantly affect any markers of sperm function assessed. Development of clinical guidance in this area is recommended to ensure HCPs deliver informed advice as lubricant use in couples trying-to-conceive may inadvertently contribute to delay in conception.

## Introduction

Vaginal lubricants are commonly used by couples to manage vaginal dryness and make intercourse more comfortable (Herbenick et al., 2010; Anderson et al., 1998). Although lubricants are of widespread availability and popularity, current regulation does not require lubricant packaging to clearly state the impact a lubricant may have on sperm function or the natural fertility process. Couples trying-to-conceive (TTC) represent a subgroup in which vaginal dryness is common (Ellington and Daughtery, 2003). A recent UK survey of over 1000 women of reproductive age actively TTC found that ∼10% of participants reported active lubricant use at the time of the survey, a proportion significantly larger than those experiencing vaginal dryness (3%) (Johnson et al., 2016). In light of such widespread use of vaginal lubricants, it is essential that patients make appropriately informed choices. However, doctor-patient communication barriers combined with limited evidence and insufficient guidance for healthcare professionals (HCPs) regarding appropriate lubricant use makes this a challenging field in which to deliver advice.

A number of studies over the last four decades have demonstrated the detrimental effects of different lubricants on sperm function (Tagatz et al., 1972; Goldenberg and White, 1975; Tulandi et al., 1982; Schoeman and Tyler, 1983; Tulandi and McInnes, 1984; Boyers et al., 1987; Frishman et al., 1992; Miller et al., 1994; Kutteh et al., 1996; Anderson et al., 1998; Agarwal et al., 2008; Vargas et al., 2011; Agarwal et al., 2013; Mowat et al., 2014; Sandhu et al., 2014). Motility, progressive motility and vitality are the most commonly assessed as markers of sperm function. Although variations in methodology and reporting make quantitative review unsuitable, the in vitro data collected is in general agreement that most commercially available lubricants negatively impact sperm function, albeit variably.

The search for a sperm-safe lubricant dates back to the late 1950’s (Kantor and Kamholz, 1957), however more recently the situation has been clarified with the United States Food and Drug Administration introducing a unique product code (‘PEB’) for personal lubricants that after sufficient testing are considered ‘gamete, fertilization and embryo compatible’ (Ellington and Clifton, 2017). Such regulation and labelling in the US does not guarantee universal HCP or patient awareness of such issues, however strives to regulate and standardise a previously unclear marketplace. However, similar regulation does not currently exist in the United Kingdom, and lubricant use in couples TTC may negatively impact chances of conception.

## Materials and Methods

### Healthcare Professional (HCP) survey

See Appendix 1 for full survey. The practices of HCPs were evaluated via a cross-sectional online survey. The survey was made using the BOS online survey tool (Available at: www.onlinesurveys.ac.uk). Participants for the survey were HCPs (doctors and nurses) working in NHS or private fertility services across Scotland. Distribution was via email and through the Scottish Fertility Network. The survey was available online for a period of approx. 8 weeks. Survey responses were anonymous at the point of collection. Thematic analysis was performed on free text response data. Descriptive statistics were used for all other data collected. Data was processed, and Figures were produced in Prism 7.

### Subjects and Ethical Approval

Semen samples were obtained from 15 healthy donors with no known fertility problems. Samples were collected by masturbation into a plastic container after 48 to 72 hours of ejaculatory abstinence. Written consent was obtained from all sperm donors and all donors were recruited in accordance with the HFEA Code of Practice (version 8) under local ethical approval (13/ES/0091) from the East of Scotland Research Ethics Service (EoSRES) REC 1.

### Sample and lubricant preparation

All samples were prepared using a discontinuous density gradient using PureCeption 40%/80% (Origio, Denmark) within 1 h of ejaculation. Density gradient prepared sperm were used as this selects motile spermatozoa that are likely to reach the site of fertilization and is consistent with similar studies referenced herein. The 80% pellet was washed, and samples were diluted to approximately 15 million sperm/mL in Quinns Sperm Washing Solution (ART-1006; Origio, Denmark). In the first experiments, motile sperm fractions from 5 different donors were incubated for 0, 10, 30 and 60 minutes at 37°C in 10% solutions of two lubricants: Pré (INGfertility, Valleyford, WA) and Aquagel (Ecolab Ltd). Aquagel Lubricating Jelly (EcoLab Ltd.) is a KMJ Class: 1 lubricant. Pré Vaginal Lubricant (ING Fertility) is a FDA gamete, fertilisation, and embryo compatible personal lubricant (Ellington and Clifton, 2017). Control samples were incubated for 0 and 60 minutes in medium alone and a 10% solution of K-Y Jelly (Reckitt Benckiser Group plc). K-Y Jelly was used as a negative control based on the numerous published accounts of its damaging effects on sperm function (Agarwal et al., 2008; Frishman et al., 1992; Tagatz et al., 1972). The 10% lubricant dilution is reflective of the concentration used in previously published studies (Agarwal et al., 2008; Anderson et al., 1998; Miller et al., 1994). The range of incubation times chosen are consistent with a previously published study suggesting that the majority of fertilising sperm migrate through the cervix within 30 minutes after ejaculation (Settlage et al., 1973). In subsequent experiments, motile sperm fractions from 5 different donors were incubated for 10 minutes at 37°C in 10% (v/v), 5% (v/v), 1% (v/v) and 0.2% (v/v) solution of Aquagel (Ecolab Ltd). 10 minutes was chosen as this was when we had previously observed changes in progressive motility following incubation with Aquagel.

### Motility analysis

Motility was evaluated at 37°C using computer-assisted-sperm analysis (SCA 5.1, Microoptic, Barcelona) attached to a microscope. For each lubricant treatment, duplicate motility assessments were done and compared. The wet-preparation was made using pre-warmed 2-cell Leja chamber (20 μm depth). At least 200 spermatozoa in at least 5 microscope fields of view were examined in each duplicate count. The acceptability of counts was determined using Table 2.1 in the WHO laboratory manual for the examination and processing of human semen and average values were only determined if the difference between the duplicates was less than the limit value. Sperm motion characteristics were assessed under a negative phase contrast objective (x200 magnification) with the system parameter settings for these analyses being 25 frames at 25 frames per second (Hz) and particle area for detection of spermatozoa head being 2–60 μm^2^. A minimum of 20 data points was used for tracking a cell. Sperm motility was classified using a four-category scheme: rapid progressive, slow progressive, non-progressive, and immotile (Barratt et al., 2011).

### Sperm Vitality

Sperm vitality was measured in 5 motile sperm fractions from 5 different donors after 60 minutes incubation with 10% Aquagel. HOS test was performed using 1 mL of a hypo-osmotic solution (0.75 mM fructose, 0.75 mM sodium citrate in distilled H_2_0; osmolality of 150 mOsm/kg) for incubation of 100 μL of prepared spermatozoa for 30 min at 37°C. 200 spermatozoa were analysed using a phase contrast microscope (x400 magnification) as curled or not curled, according to WHO 2010.

### Statistical analysis

Statistical significance for motility data was calculated using a one-way ANOVA based on the assumption that the data was approximately normally distributed. When one-way ANOVA resulted in a p-value < predefined α error (α=0.05), a post-hoc pairwise Tukey’s HSD Test was carried out to determine statistical significance between individual groups. A p-value <0.05 was considered statistically significant. As motility data for Aquagel and Pré was recorded at intermediate time points (10 and 30 minutes) which control data was not recorded at, paired t-tests compared to 0 minutes were used to determine statistical significance at these points where a p-value <0.05 was considered statistically significant.

## Results

### Healthcare professional survey

A total of n=32 responses, comprising n=19 nurses and n=12 doctors, was available for analysis. Enquiring about lubricant use in couples TTC was uncommon among respondents with 85% (n=28) of HCPs *rarely* or *never* asking about lubricant in the medical history. However, most HCPs (81% (n=26)) would not recommend use of a lubricant in couples TTC. HCPs were asked to explain why they would or would not recommend a lubricant for a couple TTC and n=19 responses were thematically analysed. Combatting vaginal dryness, uncomfortable intercourse or avoidance of intercourse were the most common reasons for recommending a lubricant and many HCPs stated that they would only recommend a lubricant for these purposes. Some respondents stated that they had never discussed this issue with patients before and one stated they had never considered lubricants as a possible cause of infertility. One respondent acknowledged the sensitive nature of this aspect of medical history taking, explaining that vaginal dryness is explored as an issue if patients identify it when asked about problems with intercourse.

Specific lubricants recommended by HCPs included Astroglide, Conceive Plus, Pre-seed, K-Y Jelly, Sliquid Oceanics Natural Lube, and ‘water-based lubricants’. Only 9% (n=3) of respondents had prescribed a lubricant for a patient in the past. Aquagel (Ecolab Ltd) was the only lubricant that was identified as having been previously prescribed. HCPs were asked if they were aware of any guidance, either local or national, addressing lubricant use in couples TTC. All respondents surveyed (n=32) stated that they were unaware of any guidance.

HCPs were asked to what extent they agreed with the following statement: ‘*Lubricants marketed for fertility patients can have an impact on sperm function*’. The majority of respondents (66% (n=21)) chose a neutral stance opting to ‘*neither agree nor disagree*’. HCPs were asked to identify the best definition, out of a possible 4, that described a non-spermicidal lubricant. Only 31% (n=10) of respondents identified the correct response: a non-spermicidal lubricant contains no drug known to kill sperm (see Figure 1).

**Figure 1.**
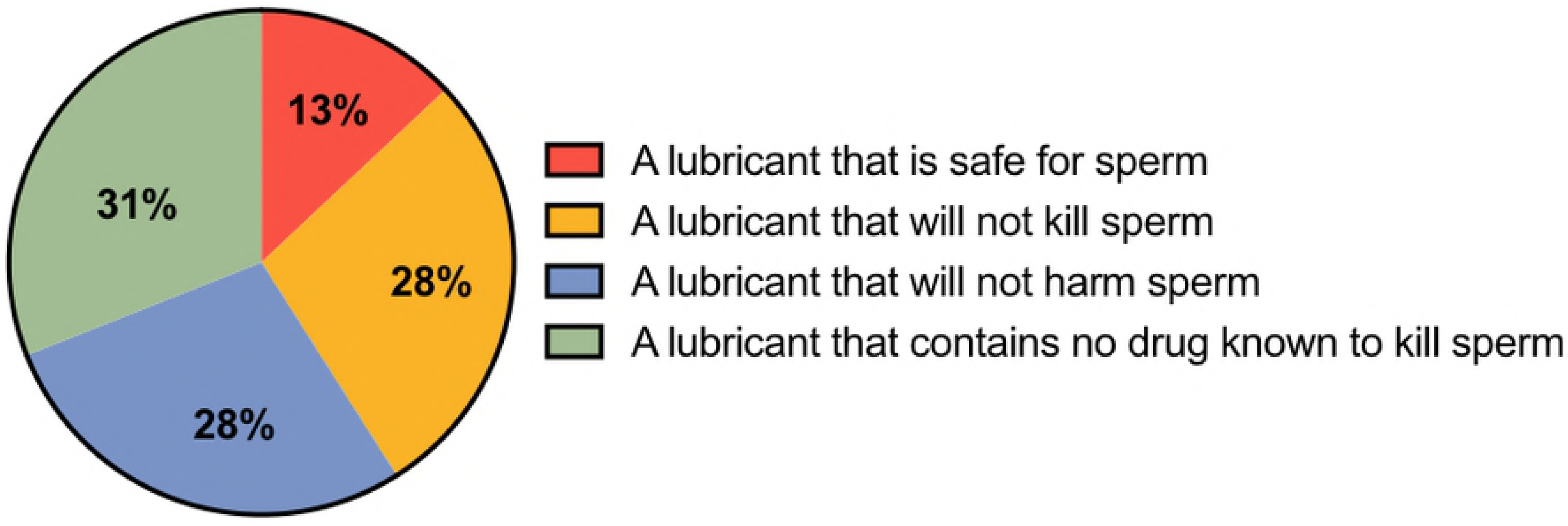
Which of the following best describes a lubricant with the classification ‘non-spermicidal’? Pie chart representing n=32 responses. The correct answer is ‘a lubricant that contains no drug known to kill sperm’.

### The in vitro effect of lubricants on sperm function

See Figure 2 for results summary. 10% (v/v) Aquagel significantly reduced progressive motility (PM) to 2% of control at 60 minutes (p<0.01). Significant reductions in PM after 10% (v/v) Aquagel exposure were also observed at 10 (p<0.01) and 30 minutes (p<0.01). Incubation with 10% (v/v) K-Y Jelly significantly reduced PM to negligible levels at 60 minutes (p<0.01). Incubation with 10% (v/v) Pré did not significantly change PM compared to control at any time assessed. Vitality analysis after 60 minutes incubation with 10% Aquagel showed no difference in vitality between 10% (v/v) Aquagel and control, suggesting PM decreases were not secondary to cell death (see Figure 2C).

**Figure 2.**
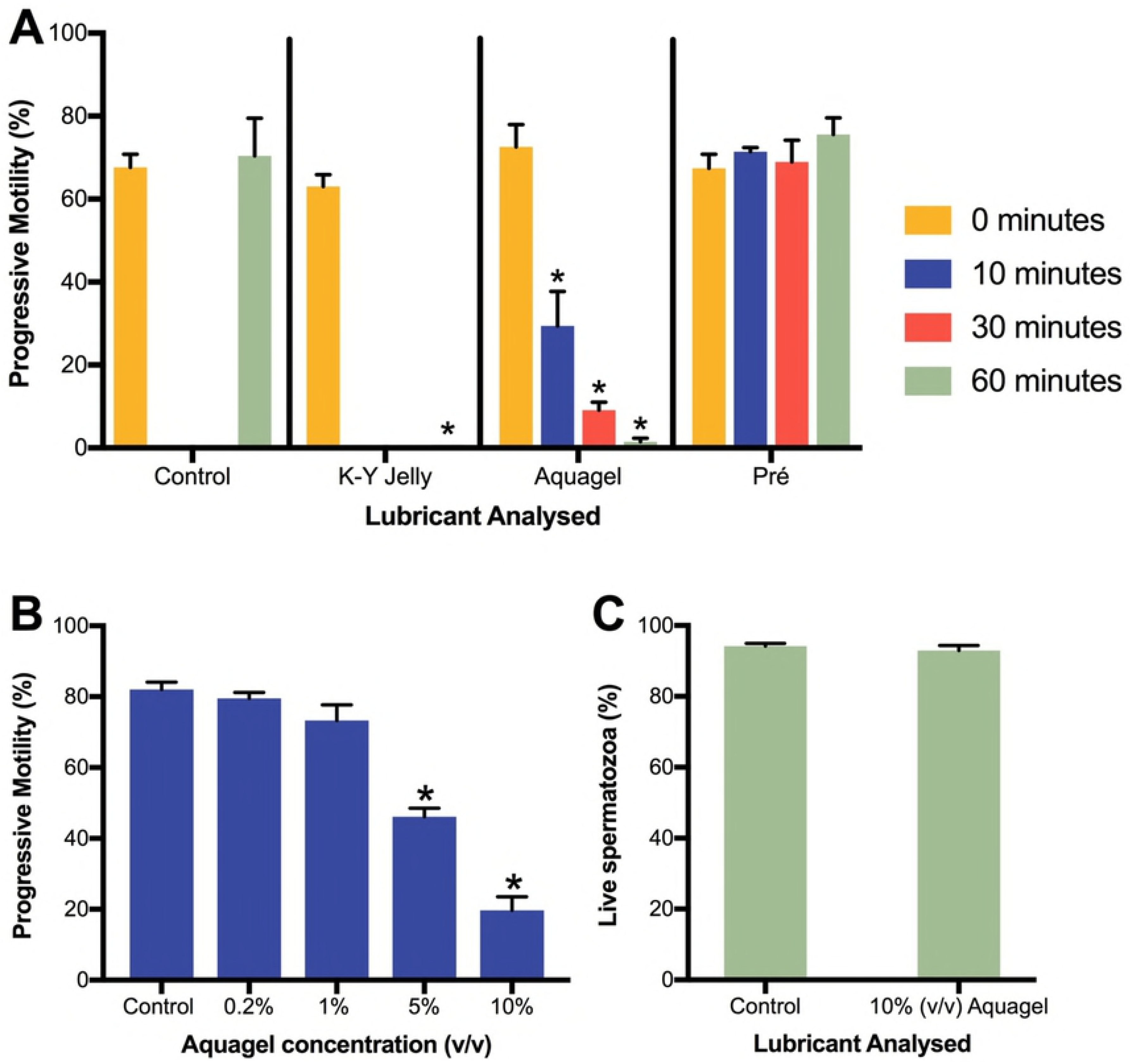
Graphs A, B and C represent data from n=5 prepared samples from 5 different donors per experiment. A represents percentage progressive motility after 60 minutes exposure to 10% (v/v) lubricant. B represents percentage progressive motility after 10 minutes exposure to Aquagel at varied concentration. C represents vitality after 60 minutes 10% (v/v) Aquagel exposure. * represents p≤0.01.

Aquagel was subsequently incubated with samples at 10% (v/v), 5% (v/v), 1% (v/v) and 0.2% (v/v), with PM assessed at 10 minutes (post-lubricant addition). 10% (v/v) Aquagel significantly reduced PM to 25.6% of control at 10 minutes (p<0.01). 5% (v/v) Aquagel significantly reduced PM to 58% of control at 10 minutes (p<0.01). No significant changes to PM were observed at 1% (v/v) or 0.2% (v/v).

## Discussion

It is unclear if HCP avoidance of lubricant use in history-taking is due to a lack of awareness or HCPs sharing the personal embarrassment, perceived unacceptability or perceived unimportance reported by patients with regard to general sexual health issues (Houge, 1988; Berman et al., 2003). A common approach across HCPs was to discuss the topic if patients identified it was an issue upon general sexual health questioning, but given that women often expect leadership from HCPs in raising sexual health issues, this approach may be inappropriate leaving many patients who are open to advice without appropriate guidance (Houge, 1988). It is therefore suggested that HCPs consider integrating querying lubricant-use in their routine history taking in the fertility setting. Additionally, educational intervention has been shown to significantly increase HCP awareness of the importance of discussing sexual health with patients and may be an appropriate method to increase lubricant-related questioning among HCPs (Blair et al., 2013).

An important distinction is that the term ‘non-spermicidal’, commonly used to classify lubricants, is a drug classification specifying that a product does not contain a spermicidal drug. As discussed by Mortimer et al., (2013), this classification does not mean a product is sperm-safe or that it will not harm or kill sperm. This misconception was common among surveyed HCPs, and is likely present among patients.

Aquagel, is a multipurpose lubricant available for NHS prescription as listed in the NHS Drug Tariff Part IX (NHS Business Services Authority, 2018). This study found that Aquagel decreased progressive motility in a time- and concentration-dependant manner, even at concentrations as low as 5% (v/v). Compared to lubricants analysed in Anderson et al., (1998), Agarwal et al., (2008) and Sandhu et al., (2014), Aquagel is one of the most sperm toxic ‘non-spermicidal’ lubricants currently available in the marketplace. In agreement with past findings (Kutteh et al., 1996; Anderson et al., 1998; Agarwal et al., 2008), K-Y Jelly was also found to be detrimental for sperm motility. Anderson et al., (1998) suggests that the high osmolality of K-Y Jelly may cause damage to tail membranes resulting in impaired motility, however the full molecular mechanism(s) of this effect awaits further investigation.

Appropriate lubricant advice is ultimately quick, easy and free and may help couples achieve natural pregnancy. Vaginal lubricants should be avoided when TTC unless specifically indicated to manage or prevent sexual dysfunction and this should be the message relayed by HCPs to patients. When indicated, Pré Vaginal Lubricant is an appropriate choice and this study supports its FDA classification as a ‘gamete, fertilization and embryo compatible’ lubricant. Aquagel should not be used or prescribed in couples TTC to manage vaginal dryness. Vaginal lubricants should be assumed to be not sperm safe and therefore not recommended or prescribed in those TTC unless robust evidence is present suggesting the contrary. It must be emphasised that although many vaginal lubricants significantly impair motility they cannot be relied upon as contraceptives and patients should be made aware of this.

Development of guidelines and patient information in this area is required to improve both HCP and patient understanding of the issues presented. Until such guidance and information is developed, clinics are encouraged to discuss these issues and develop a unified approach to lubricant recommendations to inform patient usage where appropriate.

## Acknowledgements

The authors are very grateful to Anne McConnell from the Assisted Conception Unit at Ninewells Hospital for distributing the survey to members of the Scottish Fertility Network. The authors acknowledge Dr Sarah Martins da Silva for guidelines regarding the questionnaire and Professor Christopher Barratt and Dr Vanessa Kay for their advice and comments.

## Disclosure of Interest

Authors have nothing to declare

